# Large Scale Comparative Deconvolution Analysis of the Canine and Human Osteosarcoma Tumor Microenvironment Uncovers Conserved Clinically Relevant Subtypes

**DOI:** 10.1101/2023.09.27.559797

**Authors:** Sushant Patkar, Josh Mannheimer, Stephanie Harmon, Christina Mazcko, Peter Choyke, G Tom Brown, Baris Turkbey, Amy LeBlanc, Jessica Beck

**Author notes:** co-first authors (Sushant Patkar, Josh Mannheimer).

## Abstract

Osteosarcoma is a relatively rare but aggressive cancer of the bones with a shortage of effective biomarkers. Although less common in humans, Osteosarcomas are fairly common in adult pet dogs and have been shown to share many similarities with their human analogs. In this work, we analyze bulk transcriptomic data of 213 primary and 100 metastatic Osteosarcoma samples from 210 pet dogs enrolled in nation-wide clinical trials to uncover three Tumor Microenvironment (TME)-based subtypes: Immune Enriched (IE), Immune Enriched Dense Extra-Cellular Matrix-like (IE-ECM) and Immune Desert (ID) with distinct cell type compositions, oncogenic pathway activity and chromosomal instability. Furthermore, leveraging bulk transcriptomic data of canine primary tumors and their matched metastases from different sites, we characterize how the Osteosarcoma TME evolves from primary to metastatic disease in a standard of care clinical setting and assess its overall impact on clinical outcomes of canines. Most importantly, we find that TME-based subtypes of canine Osteosarcomas are conserved in humans and predictive of progression free survival outcomes of human patients, independently of known prognostic biomarkers such as presence of metastatic disease at diagnosis and percent necrosis following chemotherapy. In summary, these results demonstrate the power of using canines to model the human Osteosarcoma TME and discover novel biomarkers for clinical translation.

## Introduction

Osteosarcoma (OS) is a rare malignancy of the bone that most commonly affects children. Despite aggressive treatment regimens that typically include additional surgery, radiotherapy, and chemotherapy, 5-year survival rates of patients with osteosarcoma have plateaued around 60-70%^1^. Hence there is a clinically unmet need for biomarkers that can stratify patients into meaningful subgroups and open avenues for precision oncology. With the absence of a clear druggable, driving molecular target in Osteosarcoma^2^, several groups have turned their attention towards the tumor microenvironment (TME) aiming to identify immune-related transcriptional signatures associated with favorable prognosis^3-11^. However, due to shortage of data, the prognostic value of most signatures has not been comprehensively evaluated in multiple external cohorts, especially in the context of clinically established prognostic features such as the stage of disease at diagnosis and percent tumor necrosis following neoadjuvant chemotherapy^12^. Furthermore, despite demonstrating immunogenic characteristics, recent immunotherapy-based clinical trials in Osteosarcoma have yielded disappointing results^13-17^, suggesting that we still lack an in depth understanding of how the primary and metastatic OS TME suppresses anti-tumor immune responses.

To better understand disease progression in Osteosarcoma, several preclinical murine models have been developed^18^. However, despite their increasing sophistication, they still fail to capture the sheer complexity of the tumor itself and a co-evolving microenvironment, inclusive of different tumor–associated stroma and immune components. Naturally occurring canine osteosarcomas, on the other hand, have strong biological and molecular similarities with human osteosarcoma^19-24^ and are estimated to be at least 10 times more common in pet dogs compared to humans^25^. Pet dogs develop tumors spontaneously in the face of an intact, educated immune system and comparable environmental exposures to humans. Hence, they are a potentially useful resource to study the Osteosarcoma TME and test the robustness and generalizability of biomarkers^11,26^.

In this work, we perform bulk transcriptomic deconvolution of 213 primary and 100 metastatic Osteosarcoma samples derived from 210 pet dogs enrolled in COTC021/022 clinical trials conducted by the Comparative Oncology Trials Consortium (COTC)^27^. This analysis uncovers distinct clinically relevant tumor microenvironment subtypes, which we validate using targeted Nanostring profiling and immunohistochemical staining of canine osteosarcoma specimens. In contrast to prior work, which has extensively analyzed the immune infiltrate of Osteosarcomas, our analysis sheds insights on the role of additional overlooked components of the Osteosarcoma TME such as the extra-cellular matrix and tumor vasculature on anti-tumor immune responses and overall disease progression. Furthermore, utilizing data of primary and matched metastatic tumor samples from canines, we chart how the Osteosarcoma TME evolves from primary to metastatic disease in a clinical setting and assess its impact on overall survival outcomes of canines receiving standard of care chemotherapy treatment. Most importantly, with the help of machine learning, we build a TME subtype-based risk stratification model and show that it generalizes from canine to multiple human Osteosarcoma datasets. Unlike previous studies, we comprehensively benchmark our proposed TME subtype-based risk stratification model against varying clinical features of both canine and human patients and demonstrate its prognostic value independent of previously established prognostic factors. Overall, this work highlights the tremendous potential of utilizing canines to derive generalizable biomarkers for human Osteosarcoma and inform personalized immunotherapeutic treatment strategies.

## Results

### The tumor microenvironment landscape of canine Osteosarcomas

To comprehensively characterize different cellular components of the Osteosarcoma TME, we first performed bulk transcriptomic deconvolution of 198 canine tumor samples (190 treatment naïve primary, 8 metastatic) with the help of MCP-counter^28^: an enrichment-based deconvolution algorithm which estimates the relative abundance of major cell type lineages expected to be present in each sample (Figure 1A, See more details in Methods). All 198 samples were profiled using bulk RNA Sequencing and obtained from a cohort of 186 dogs enrolled in the COTC021/022 trials (*Discovery Cohort*). Consensus clustering analysis^29^ of deconvolved transcriptomic profiles from the discovery cohort revealed three distinct TME subtypes (Figure 1B, Supplementary Figure 1A-D): (i) Immune enriched (IE), which consists of tumors that are enriched for Cytotoxic T and NK cell populations, (ii) Immune enriched dense extra-cellular matrix like (IE-ECM) which consists of tumors with a diverse milieu of infiltrating immune cells and in addition are enriched for endothelial and ECM/Stromal subpopulations (iii) Immune Desert (ID), which are predominantly cold tumors with low levels of immune infiltrate. The observed subtypes are independently recapitulated by Nano string data of 23 primary and 92 metastatic samples biopsied from 40 dogs at necropsy (*Validation Cohort*, Figure 1C, Supplementary Figure 2), and in addition, by another state-of-the-art deconvolution algorithm, xCell^30^ which considers a wider range of cell types and cell states (Supplementary Fig 3A-D). On performing subtype-specific differential expression and functional enrichment analysis (See Methods), we observe that the top 5 hallmark pathways enriched in the IE subtype are ALLOGRAFT_REJECTION, INTERFERON_ALPHA_RESPONSE, INTERFERON_GAMMA_RESPONSE, INFLAMMATORY_RESPONSE and TNFA_SIGNALING_VIA_NFKB. The top 5 hallmark pathways enriched in the IE-ECM subtype are: IL6-JAK-STAT3_SIGNALING, COAGULATION, INFLAMMATORY_RESPONSE, KRAS_SIGNALING_UP, and TNFA_SIGNALING_VIA_NFKB, whereas the ID subtype shows a complete lack of an immune response with the following top 5 enriched hallmark pathways: E2F_TARGETS, MYC_TARGETS_V1, G2M_CHECKPOINT, MYC_TARGETS_V2 and OXIDATIVE_PHOSPHORYLATION (Supplementary Tables 1-3). Given that chromosomal instability (CIN) is also a characteristic feature of Osteosarcomas^31^, we additionally estimated the level of CIN in each sample using a previously defined gene expression signature of CIN^32^. Interestingly, we observe significant differences in estimated levels of CIN across subtypes with the ID subtype having the highest estimated levels of CIN, closely followed by the IE-ECM and IE subtypes (Figure 1E).

**Figure 1.**
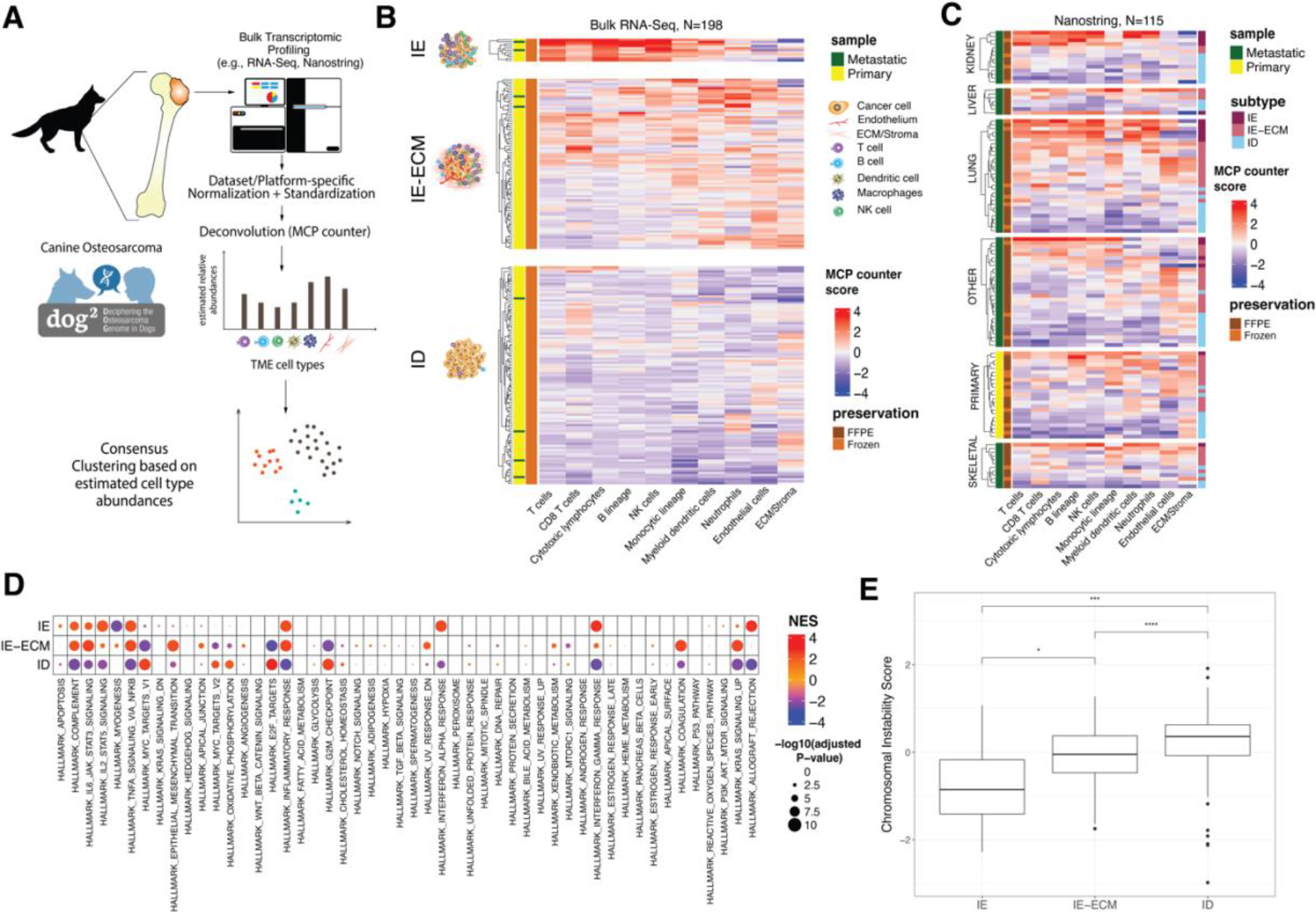
Overview of the TME landscape of canine Osteosarcomas. **A:** Overview the computational pipeline for characterizing the TME landscape of canine Osteosarcomas. **B-C**: Heatmaps depicting the MCP counter estimated relative abundances of each cell type in each sample and corresponding cluster memberships for the Discovery Cohort (B) and the Validation cohort (C). **D:** Bubble plot depicting subtype-specific gene set enrichment analysis results. NES: Normalized Enrichment Score. **E:** Boxplot depicting the Chromosomal Instability Score of each subtype. The Chromosomal Instability Score is defined as the average normalized expression of genes part of the Chromosomal Instability Signature^32^. Statistical significance between groups was determined by the non-parametric Wilcoxon rank sum test (*: p-value < 0.05, ***: p-value < 0.001, ****: p-value < 0.0001)

The IE subtype, which is characterized by significantly high levels cytotoxic T/NK cells constitutes a minority of all immune infiltrated tumors (N=11 Discovery cohort, N=17 Nanostring validation cohort) and has significantly higher expression levels of immune checkpoint genes: PD1, PDL1 CTLA4, LAG3, TIM3 and TIGIT in comparison to other subtypes (See Supplementary Figure 4A, B). The IE-ECM subtype (N=82 discovery cohort, N=54 Nanostring validation cohort) constitutes a majority of the immune infiltrated tumors and is uniquely characterized by relatively high expression of ECM, endothelial cell markers, and in addition, genes involved in upregulation of oncogenic KRAS signaling. These results suggests that Osteosarcomas frequently remodel the TME in response to host immunosurveillance via activation of specific oncogenic pathways^33^. To further validate these observations, we performed immunohistochemical staining of matched tumor samples from dogs with highest (top 10) and lowest (bottom 10) endothelial and stromal gene expression levels. Figure 2A depicts the estimated number of blood vessels in the primary TME of cases that were reviewed by a veterinary pathologist. The number of blood vessels was manually estimated from 10 randomly chosen high powered magnification fields per case. Similarly, Figure 2B depicts the pathologist estimated percentage tissue area containing tumor cells, osteoid (dense extra-cellular bone matrix) and Collagen I-III. Despite challenges associated with IHC staining and manual quantification of stromal and endothelial features from visual assessment, we observe noticeable histologic differences in the TME of tumors with high or low endothelial and stromal gene expression. The most notable difference is in the level of Osteoid production, which is elevated in tumors with high expression of ECM markers (Figure 2C) in comparison to tumors with the low expression (Figure 2D)

**Figure 2.**
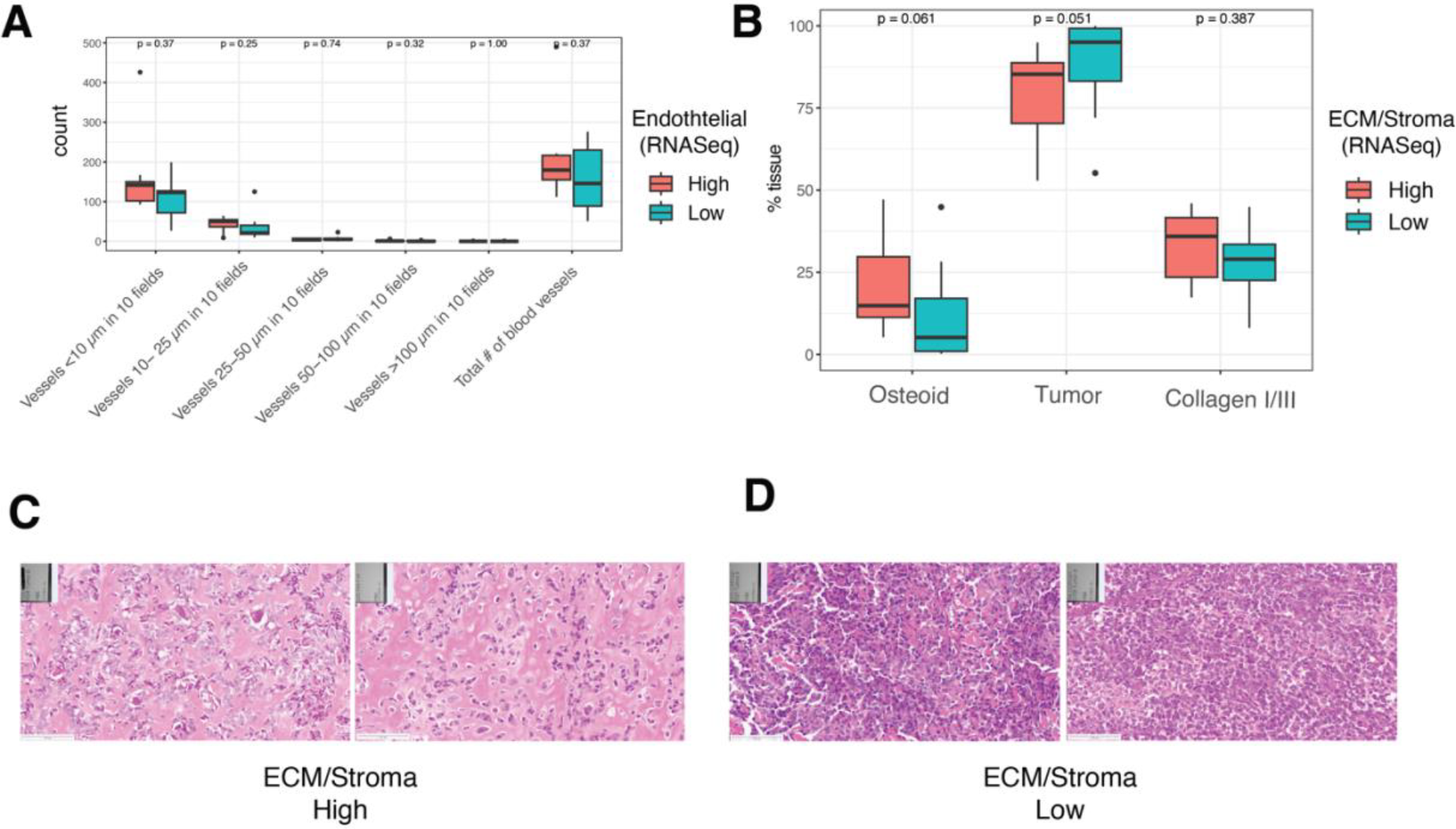
Transcriptional levels of endothelial and stromal markers are associated with endothelial and stromal features viewed on histology. **A:** number of blood vessels estimated from visual examination of pathologist and grouped by their diameter (on X axis) in cases inferred to have high endothelial expression vs low endothelial expression from transcriptomic deconvolution analysis. **B:** percent tissue area covered by Osteoid, tumor cells and Collagen I-III based on visual examination of histologic slides of tumors predicted to have high expression of ECM/Stroma markers and likewise low expression from bulk transcriptomic deconvolution. **C -D:** Two representative 20x magnification views of tumors from cases inferred to have highest expression of ECM/Stromal markers (C) and lowest expression

### Primary TME landscape and clinical outcomes of canine patients enrolled in COTC021/022

We next investigated if the TME subtypes of the primary tumor are predictive of clinical outcomes of dogs enrolled in the COTC021/022 clinical trials. All canines enrolled in these trials were metastasis-free at the time of diagnosis with most dogs eventually developing metastatic disease during the follow-up period. See Methods for additional details on sample curation and Supplementary Table 4 for all pertaining clinical metadata. Remarkably, the inferred TME subtypes of the primary tumor strongly stratify the overall survival and disease-free intervals of canines with the IE subtype having most favorable prognosis, closely followed by the IE-ECM subtype, and finally the ID subtype having the worst prognosis (Figure 3A, B; overall survival log-rank test p-value: 0.0012, DFI log-rank test p-value: 0.0017). We additionally fit a multi-variate Cox proportional hazards regression model to assess the predictive value of the primary TME subtype in the context of other clinically measured variables such as age, weight, primary tumor location (proximal vs non-proximal to humerus), gender, alkaline phosphatase (ALP) levels and clinical trial treatment arm (standard of care chemotherapy with or without Sirolimus). Overall, the primary TME subtype has a significant effect on disease free intervals even after accounting for other clinical factors (Figure 3C, D). This suggests that the primary TME composition could serve as an independent prognostic biomarker for metastatic disease progression and stratification of patient risk.

**Figure 3.**
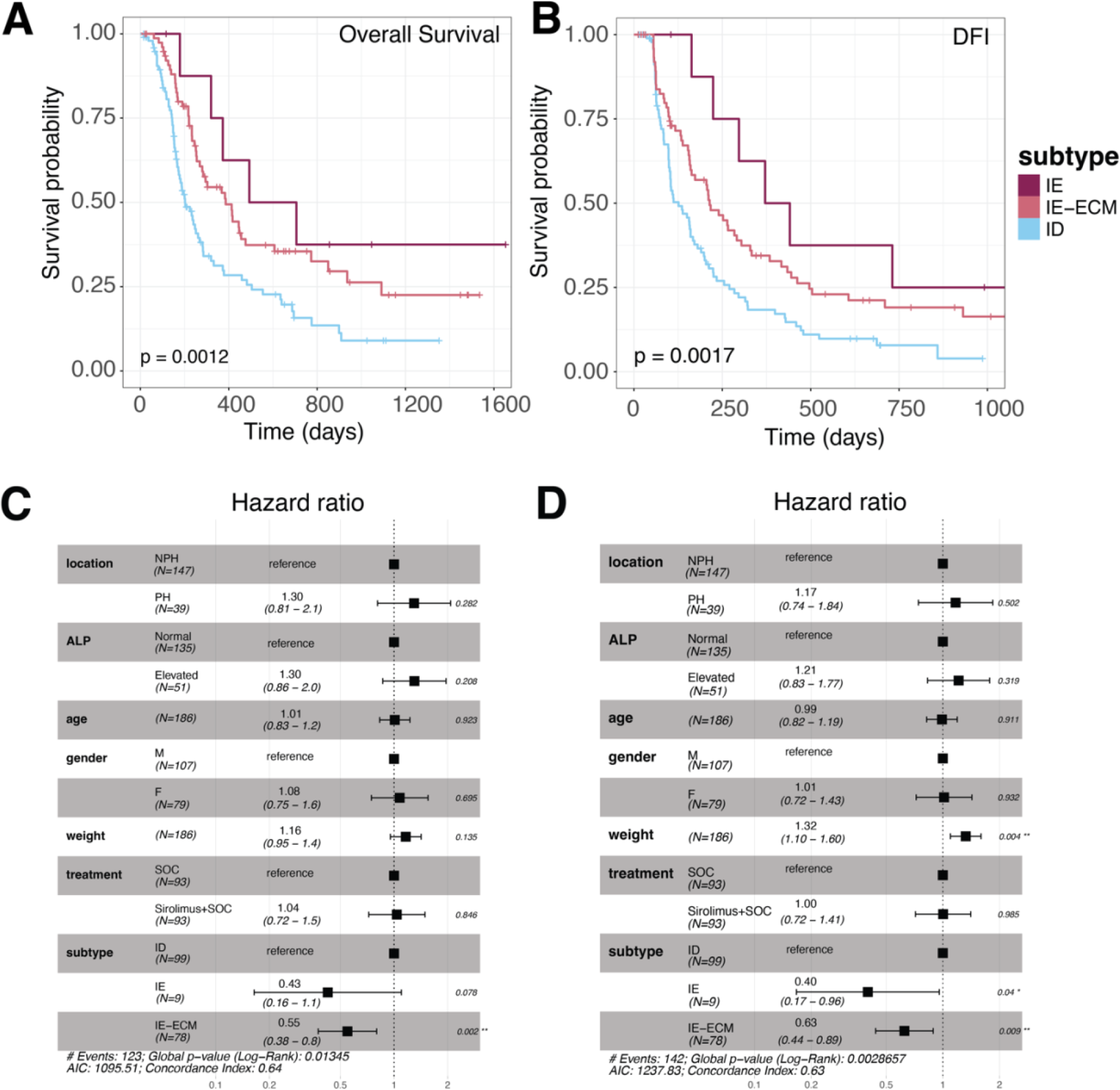
Relationship between primary TME subtype and clinical outcomes in DOG2. **A-B:** Kaplan-Meier survival plots for overall survival (A) and disease-free intervals (B) of canines enrolled in COTC021/022 trials and stratified by the primary tumor TME subtype. Statistical significance of differences between survival trajectories was determined by the log rank test **C-D:** Forrest plots, depicting estimated hazard ratios of clinical features and TME subtype from cox proportional hazards regression analysis. (*: p-value < 0.05, **: p-value < 0.01)

### Evolution of the Osteosarcoma tumor microenvironment from primary to metastatic disease in a standard of care clinical setting

Despite the fact that Osteosarcoma frequently metastasizes to distinct organs, very little is known about its metastatic TME landscape and how it relates with primary tumors of patients receiving standard of care treatment. This is mainly due challenges associated curating tumor samples from human patients^1,2^. However, fewer barriers exist for collecting and analyzing tumor samples from dogs. This makes canine Osteosarcomas a natural model for studying the evolution of these relatively rare tumors from primary to metastatic disease in a clinical setting. When comparing the TME composition of primary tumors to their matched metastases from a variety of geographic locations, no distinct patterns emerged suggesting that the observed TME subtypes are both conserved and switched as tumors evolve from primary to metastatic disease (Figure 4A). However, despite the heterogeneity of the metastatic TME, we observed that metastatic samples from canines with more favorable clinical outcomes (overall survival time > median of cohort) had significantly higher levels of cytotoxic T cell populations compared to canines with less favorable outcomes (overall survival time < median), irrespective of metastatic site (Fig 4B). Whereas metastatic samples from canines with less favorable outcomes had significantly higher levels of endothelial and stromal populations compared to canines with more favorable clinical outcomes (Fig 4B). These results suggest that besides the primary TME composition, an increased presence of cytotoxic T cells and decreased presence of tumor-associated stroma and endothelial populations at different metastatic sites positively impacts overall survival of canine patients.

**Figure 4.**
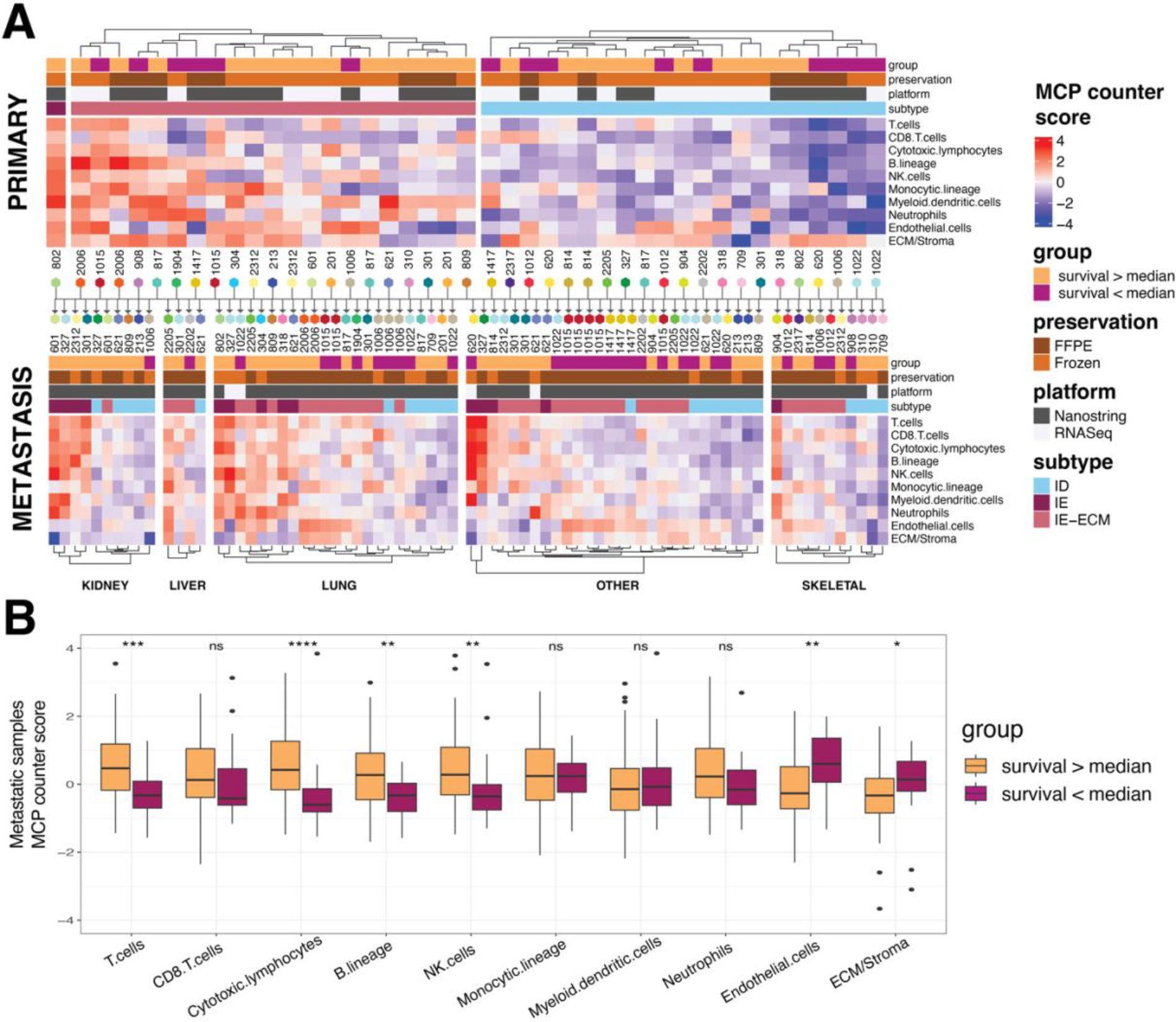
Evolution of the Osteosarcoma TME from primary to metastatic disease. **A**: Heatmap depicting the MCP counter-estimated relative cell type abundances in primary and matched metastatic tumors from 28 cases that were sampled at diagnosis and at necropsy. **B:** Boxplots depicting the MCP counter-estimated relative cell type abundances in metastatic tumor samples from different sites and grouped by overall survival duration. Statistical significance between overall survival groups was estimated using the wilcox rank sum test (p-value < 0.05: *, p-value < 0.01: **, p-value < 0.001: ***, p-value < 0.0001: ****)

### Translation of biomarkers from canines to humans using machine learning

Although canine Osteosarcomas have been shown to have strong similarities with human Osteosarcomas, it is yet unknown whether biomarkers derived from the analysis of canine data can reliably stratify clinical outcomes of human patients. To test this hypothesis, we trained a simple Naïve Bayes classifier^34^ to predict the TME subtype from the estimated TME composition of each canine tumor sample (Figure 5A, See Methods for more details). The TME subtype classifier, on average, achieves a 10-fold cross-validation accuracy of 91.5%, CI: [90%-93%] and kappa score of 0.872, CI: [0.85-0.89] when trained and validated on canine tumor data. We then applied the trained naïve bayes classifier on seven independent publicly available Human Osteosarcoma bulk gene expression datasets (N=394) and in addition one single cell dataset (N=11) pre-processed using MCP counter to predict the TME subtype of each human sample^11,35-41^. Majority of samples curated and analyzed in this study represent pre-chemotherapy diagnostic samples (N=383 of 405). Detailed clinical metadata for each dataset evaluated is provided in Supplementary table 5. Figure 6B shows a heatmap of MCP counter-estimated relative cell type abundances in human Osteosarcoma samples along with their model-predicted TME subtype. Overall, the estimated cell type composition, chromosomal instability, and immune checkpoint gene expression levels across inferred TME subtypes of human Osteosarcoma samples is similar to what was observed in canines (Figure 5B, C, Supplementary Figure 5A,B, Supplementary Figure 6). Furthermore, we applied the classifier on merged single cell expression data of 11 human Osteosarcoma patients profiled using single cell RNA Seq. Supplementary Figure 7 A,B summarizes the TME composition of 11 tumor samples based on previous single cell analysis^41^. Despite the low sample size, we observe that tumor samples classified as the IE subtype (N=1) are uniquely characterized by elevated levels of cytotoxic CD4+ T cell, CD8+ T cell and NK cell populations at the single cell resolution, and in addition antigen presenting Dendritic cell subpopulations (CD14+/CD163+, CD1C+ DCs), which have been previously linked with favorable responses to immunotherapies^42^. Samples classified as the IE-ECM subtype (N=4) are uniquely characterized by elevated levels of mesenchymal stem cell and malignant cell populations of Osteoblastic and chondroblastic lineage (Osteoblastic_3, Osteoblastic_4, Osteoblastic_5, Osteoblastic_6, Chondro_proli, Chondro_hyper1, Chondro_hyper2, Chondro_trans). These populations were shown to express diverse transcriptional programs involved in an inflammatory response, and in addition, angiogenesis, myogenesis, oxidative phosorylation, hypoxia, MYC, TGF-beta, TP53 and KRAS signaling^41^. Lastly, the ID subtype (N=6) was uniquely characterized by elevated levels of a highly proliferative subpopulation of Osteoblastic cancer cells (Osteoblastic_1 and Osteoblastic_2).

**Figure 5.**
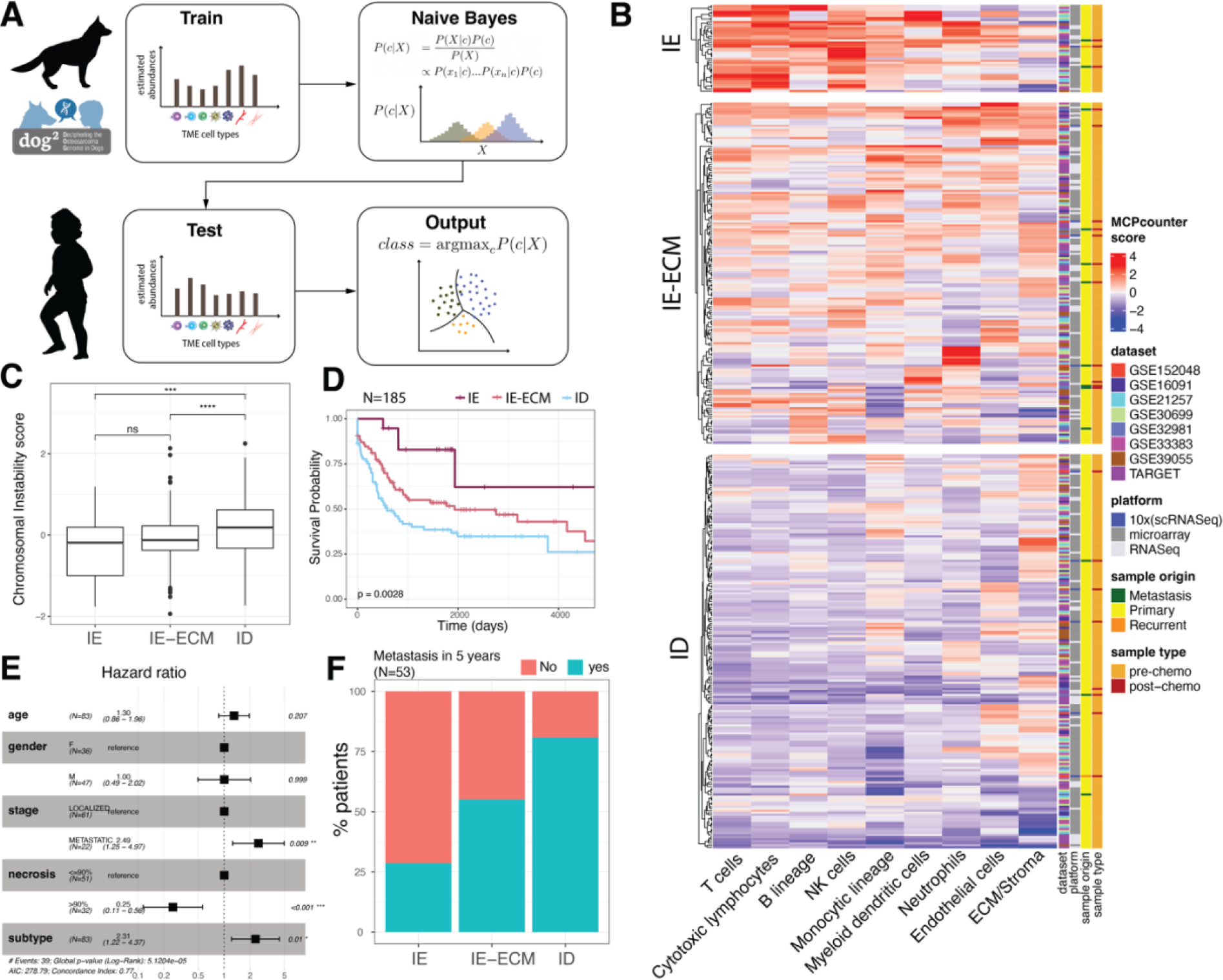
Relationship between primary TME subtype and clinical outcomes in Humans. **A:** Overview of the machine learning pipeline to predict TME subtype of canine and human Osteosarcomas. **B:** Heatmap depicting the MCP counter-estimated relative cell type abundances and predicted TME subtypes of 405 human Osteosarcoma samples based on training on canine tumor data. **C:** Boxplot depicting the Chromosomal Instability Score of each subtype. The Chromosomal Instability Score is defined as the average normalized expression of genes part of the Chromosomal Instability Signature^32^. Statistical significance between groups was determined by the non-parametric Wilcoxon rank sum test (***: p-value < 0.001, ****: p-value < 0.0001). **D:** Kaplan-Meier plots depicting the progression free survival of human Osteosarcoma patients grouped by their primary tumor TME subtype. Statistical significance of differences between survival trajectories was determined by the log rank test. **E:** Estimated hazard ratios of age, gender, disease stage (localized vs metastatic), necrosis (> 90%, <=90%) and TME subtype (IE subtype: 1 or low risk, IE-ECM subtype: 2 or intermediate risk, ID subtype: 3 or high risk) based on cox-proportional hazards regression analysis of 83 patients spanning three datasets: TARGET^35^, GSE39055^36^, GSE32981^38^. **F:** Stacked bar plots depicting the percentage of patients developing metastasis in 5 years from diagnosis in each subtype. Based on 53 of 84 patients from GSE33383^11^ with matched gene expression and clinical metadata

Next, when comparing the TME subtype of primary tumor samples with matched clinical outcomes of patients, we observe that model predicted TME subtypes strongly stratify progression free survival of human patients (N=185) in a manner similar to what was observed in canines (Figure 5D). Importantly, we find that the primary tumor TME subtype when considered as a prognostic indicator (i.e IE subtype = 1 or low risk, IE-ECM subtype = 2 or intermediate risk, ID subtype = 3 or high risk) has a significant effect on progression free survival (HR: 2.31[1.22 – 4.37], p-value: 0.01) even in the context of other clinically measured prognostic variables such as disease stage at diagnosis (HR: 2.49 [1.25 – 4.97], p-value: 0.009) and percentage necrosis post neoadjuvant chemotherapy (HR: 0.25 [0.113 – 0.56], p-value < 0.001), the current gold standard for evaluating response to standard of care treatment in humans^12^ (Figure 5E). Furthermore, in an independent dataset of patients with primary tumor samples (GSE33383, N=53), the inferred TME subtypes are strongly predictive of developing metastasis within 5 years (Figure 5F; chi-squared test p-value: 0.021). In summary, these retrospective analyses suggest that canines could serve as a useful real-world clinical model for analyzing the OS TME and translation of biomarkers.

## Discussion

In this work we performed an unsupervised analysis of the tumor microenvironment of canine and human Osteosarcoma patients to uncover three distinct TME subtypes that are conserved across both species and validated independently using Nanostring and IHC data from canines. To the best of our knowledge, this study represents the largest and most comprehensive comparative analysis of the Osteosarcoma TME in both canines and humans spanning multiple datasets. Importantly, in contrast to prior works we show that the uncovered TME subtypes are robustly predictive of progression free survival outcomes of both canine and human patients, independent of previously studied prognostic factors such as tumor location and serum alkaline phosphatase levels in the case of dogs, and tumor stage and percentage necrosis in the case of humans. Furthermore, we observe a strong correlation between the TME subtype and chromosomal instability of Osteosarcomas in both canines and humans. Chromosomal instability is a hallmark feature of multiple cancers, including Osteosarcomas and is believed to be one of the central players in accelerating tumor evolution, drug resistance and metastasis^43,44^. In Osteosarcomas, chromosomal instability has been shown to arise from global hypomethylation of DNA^45^. However, the effect of chromosomal instability on the surrounding TME is complex and an ongoing area of investigation. Although chromosomal instability can trigger innate immune-mediated clearance of cancer cells via the activation of cGAS-STING pathway^44^, several immune escape mechanisms have been proposed^46^. One of them being silencing of genes encoding type 1 interferons^47-49^ downstream of the cGAS-STING pathway. In both canine and human Osteosarcoma samples analyzed in this study, expression of type 1 interferon response genes is lowest in the ID subtype, the one with highest chromosomal instability (See Supplementary Figure 8 A, B). This is also recapitulated at the single cell level, where we see tumor samples belonging to the ID subtype have elevated levels of proliferative cancer cell subpopulations (Osteoblastic_1 and Osteoblastic_2; Supplementary Figure 7B) previously shown to lack expression of type 1 interferon response genes^41^.

When comparing against recent literature, we find that the TME subtypes uncovered in this work bear closest resemblance to three of four recently identified TME subtypes that were shown to be conserved across several different adult cancer types and predictive of responses to immune checkpoint blockade therapy^50^. The IE subtype uncovered in this work recapitulates the IE/Non-Fibrotic subtype identified across several other cancer types and was shown to have the most favorable responses to immune checkpoint blockade therapies irrespective of tissue of tumor origin^50^. In both canine and human Osteosarcomas, only a small fraction of tumors (∼10%) belong to this subtype. Majority of immune infiltrated tumors in Osteosarcoma in fact belong to the IE-ECM subtype, which corresponds to the immuno-suppressive IE/Fibrotic subtype from pan cancer analysis of Bagaev et al^50^. These findings potentially explain the lack of efficacy of immune checkpoint inhibitors in recent clinical trials of Osteosarcoma despite the presence of an immune infiltrate^13-15^ implying that targeting cellular populations governing ECM production in the Osteosarcoma TME in combination with immunotherapy may be an effective strategy to improve patient outcomes^51^. The ID subtype in Osteosarcoma recapitulates the Depleted subtype observed in several other cancer types and is linked with intrinsic resistance to immunotherapy^52^. The high levels of chromosomal instability observed in this subtype coupled with low type 1 interferon gene expression suggest that these tumors could potentially be susceptible to combination immunotherapy treatments that reverse silencing of type 1 interferon genes. However, sustained type 1 interferon signaling can also promote immune suppression in the TME and resistance to immune checkpoint inhibition^53^. Hence, additional research is required to precisely determine in which context type 1 interferon expression can enhance immune responses of patients with immunologically cold tumors.

Despite the merits of this study, there are some limitations that should be noted and improved upon in future work. First, the issue of pet owners’ decisions for euthanasia can confound accurate estimation of the effect of TME subtype on overall survival outcomes. This complicates comparative assessment of prognostic biomarkers in canines and humans. However, in contrast to previous comparative studies of Osteosarcomas, which focus on overall survival^11,54^, we focused on disease free intervals and progression free survival outcomes of canines and humans, respectively. This alleviates biases associated with the wishes of individual pet owners and enhances the translational value of the COTC021/22 clinical trial dataset. However, additional prospective validation is still required to fully realize the translational potential of the canine-derived biomarkers uncovered in this work. Second, majority of the data analyzed in this study was derived from bulk transcriptomic profiling of tumor biopsies masking the underlying intratumor heterogeneity present in each sample. Further comparative analysis of canine and human Osteosarcomas based on spatial transcriptomics sequencing of tumors is likely to provide more detailed insights into how malignant Osteosarcoma cells interact with their surrounding TME to facilitate immunotherapy resistance and metastasis.

## Materials and Methods

### Curation and pre-processing of canine osteosarcoma bulk RNASeq data (Discovery Cohort)

Canine Osteosarcoma tumor samples were curated from a multisite clinical trial of 324 dogs^27^. Prior to enrollment, patient tumor biopsies were examined and diagnosed as osteosarcoma by veterinary anatomic pathologists at Comparative Oncology Trials Consortium (COTC) institutions (*https://ccr.cancer.gov/comparative-oncology-program/consortium*, last accessed May 13, 2022). Following diagnosis, additional treatment-naïve tumor tissue was collected at the time of surgical limb amputation by COTC investigators as a part of the trial schema. Dogs were randomized to receive either standard of care or standard of care + adjuvant sirolimus (rapamycin) therapy. Frozen tumor samples (190 primary, 8 metastatic) from 186 out of 324 dogs were moved forward for bulk RNA sequencing analysis after completing sample QA/QC analysis and medical record review. All samples forwarded for mRNA sequencing had a RIN > 8 and a total RNA quantity > 100 ng. RNA was isolated from canine frozen tumor tissue in RNAlater using Qiagen Allprep DNA/RNA Mini Kit (Cat#80204). The total RNA quality and quantity was assessed using Nanodrop 8000 (Thermofisher) and Agilent 4200 Tapestation with RNA Screen Tape (Cat# 5067-5576) and RNA Screen Tape sample Buffer (Cat#5067-5577).

Between 100ng to 1ug of total RNA was used as the input for the RNA sequencing libraries. Libraries were generated using the TruSeq Stranded mRNA library kit (Illumina) according the to the manufacturers protocol. The libraries were pooled and sequenced on NovaSeq S1 using a 2×150 cycle kit. The HiSeq Real Time Analysis software (RTA v.3.4.4) was used for processing raw data files. The Illumina bcl2fastq2.17 was used to demultiplex and convert binary base calls and qualities to fastq format. The samples had 44 to 61 million pass filter reads with more than 91% of bases above the quality score of Q30. Reads of the samples were trimmed for adapters and low-quality bases using Cutadapt. The trimmed reads were mapped to the CanFam4 reference genome (GSD_1.0^55^ from NCBI) using STAR aligner (version 2.7.0f) with two-pass alignment option. RSEM (version 1.3.1) was used for gene and transcript quantification based on the CanFam4 GTF file. The average mapping rate of all samples was 83% with unique alignment above 66%. There were 13.13-26.26% unmapped reads. The mapping statistics were calculated using Picard software. The samples had between 0.01-0.76% ribosomal bases. Percent coding bases were between 58-71%. Percent UTR bases were 10-16%, and mRNA bases were between 75-82% for all the samples. Library complexity was measured in terms of unique fragments in the mapped reads using Picard’s “ MarkDuplicate” utility. The samples had 48-78% non-duplicate reads. In addition, the gene expression quantification analysis was performed for all samples using STAR/RSEM tools (version 1.3.1). The raw read counts for each gene were used for downstream differential expression analysis, using edgeR (v3.40.2) and gene set enrichment analysis using the clusterProfiler package (v4.6.2). The normalized read counts in log Transcripts Per Million (log2(TPM+1)) scale were utilized for downstream deconvolution analysis using MCP counter and xCell.

### Curation and pre-processing of canine osteosarcoma nanostring data (Validation Cohort)

Approximately 25% (83/324) of the total dogs enrolled in COTC021/022^27^ underwent a postmortem examination^56^. During necropsy, additional frozen and formalin-fixed samples were collected from multiple tissues including any metastases. Formalin-fixed tissue samples were collected in 10% neutral buffered formalin, decalcified using 12% EDTA (pH 7.2), and embedded in paraffin. Diagnosis of osteosarcoma metastases was based on the finalized necropsy report which included both gross and histologic examination of canine tissues by veterinary anatomic pathologists at the COTC institutions. Histologic analysis of each sample was performed to identify regions of high tumor content. Samples with <50% viable tumor tissue were annotated and scraped. Of 83 dogs, necropsy samples from 40 dogs were deemed viable and moved forward for additional Nanostring analysis. For the majority of FFPE blocks, RNA was extracted from scrolls. RNA isolation from FFPE tissue (14 primary tumors, 78 metastases) was performed by the Molecular Histopathology Laboratory (NCI). In addition, RNA was extracted from 23 frozen samples (9 primary, 14 metastases). RNA quality was assessed using the Agilent TapeStation to identify samples for which at least 50% of the RNA fragments were greater than 200 nucleotides in length (DV200). RNA was profiled using the Canine IO Panel and analyzed with the nCounter System (NanoString Technologies).

### Curation and preprocessing of FFPE slides for Immunohistochemistry validation

A subset of primary tumor FFPE samples from COTC021/022 were identified based on their disparate expression of endothelial or fibroblast-associated genes from bulk RNA Sequencing. Following histologic review to identify sections with high tumor content, primary tumor cases were selected and categorized as endothelial high (n=10), endothelial low (n=10), fibroblast high (n=10), and fibroblast low (n=10) based on transcriptomic data analysis. The 20 endothelial FFPE samples were labelled for Factor VIII-related antigen (F8ra) by the Histology Laboratory at the University of Georgia’s College of Veterinary Medicine. The 20 fibroblast FFPE samples were labelled for Collagen Types I/III (Abcam; Ab24137) by VitroVivo. Immunohistochemical results were analyzed in the HALO (indica labs) digital pathology software suite.

### Curation and pre-processing of publicly available human Osteosarcoma datasets

A total of 8 publicly available human Osteosarcoma bulk gene expression and single cell RNASeq datasets were analyzed in this study: TARGET(N=87)^57^, GSE39055(N=37)^36^, GSE33383(N=84)^37^, GSE32981(N=23)^38^, GSE30699 (N=76)^39^, GSE21257 (N=53)^40^, GSE16091 (N=34)^11^, GSE152048 (N=11)^41^. Pre-processed bulk RNA sequencing data of Human Osteosarcomas from the TARGET study was obtained from the UCSC xena browser (www.xenabrowser.net/datasets). Specifically, gene expression was quantified from RNA sequencing data using STAR (2.7.10b) and gencode v22 human genome annotation and normalized to log2(TPM+1) scale. For more details on the expression quantification pipeline, see https://docs.gdc.cancer.gov/Data/Bioinformatics_Pipelines/Expression_mRNA_Pipeline/.

Normalized microarray gene expression datasets were curated based on an initial search on Gene Expression Omnibus (GEO) database using the following keywords: “ Osteosarcoma”, “ biopsy”, and “ homo sapiens”. Eventually all datasets with sample size > 20 or matched clinical metadata documenting progression free survival outcomes were selected for further analysis. Besides bulk gene expression, pre-processed Single cell RNASeq UMI count data of 11 human Osteosarcomas was obtained from the GEO database using the following study accession ID: GSE152048^41^. Single cell RNASeq UMI count data was further filtered following the procedure described in the corresponding study to remove cells that were potentially doublets or contained < 300 expressed genes. Filtered single cell gene expression profiles from each sample were then averaged and normalized to log2(TPM+1) scale to generate one bulk gene expression profile per sample.

### Deconvolution analysis

To facilitate comparative deconvolution analysis across multiple datasets while controlling for differences associated with different gene expression platforms and sample preservation techniques, all pre-processed gene expression data was further standardized to z-scores based on dataset-specific mean and standard deviation of the normalized expression of each gene in each dataset^50^. The standardized gene expression values were then provided as input to MCP counter, a gene set enrichment-based deconvolution algorithm that quantifies the relative abundance of each cell type/lineage in terms of the average expression of a set of previously defined marker genes uniquely expressed in each cell type/lineage^28^. MCP counter by default estimates the relative cell type abundances of the following cell types/lineages expected to be present in the tumor microenvironment: CD8+ T cells, CD4+ T cells, Cytotoxic Lymphocytes, B cells, NK cells, Monocytic lineage, Myeloid lineage, Neutrophils, Endothelial cells and ECM/Stroma/Fibroblasts. The resulting relative abundances of each cell type/lineage correspond to the *TME profile* of each sample. For additional validation using a more extended reference cell atlas, we performed deconvolution using xCell^30^, another highly cited state-of-the-art gene set enrichment-based deconvolution algorithm which defines marker genes for 64 distinct cell types/cell states expected to be present in normal or cancerous tissue samples. Besides MCP counter and xCell, there are several other more sophisticated deconvolution algorithms published in literature^58-63^. However, we report our results based on MCP counter for the following reasons: i) simplicity and flexibility of implementation across various gene expression platforms compared to other approaches ii) built on a robust set of marker genes that are uniquely expressed in each cell type, mitigating issues associated with spill-over of markers from closely related cell types compared to other approaches^64^ iii) in contrast to other approaches approaches, which aim to estimate fractional cell type abundances that add up to 1, MCP counter estimates relative abundances of cell types independently of one another. This allows unbiased application of this approach to cancer types/tissues such as Osteosarcomas, where the presence or absence of specific cell types is not precisely defined^64^.

### Consensus clustering and machine learning analysis

Following deconvolution analysis with MCP counter, the standardized TME profiles of each sample from the discovery cohort were analyzed using the consensusClusterPlus package^29^ (v1.62.0). The following parameters were specified as input for the unsupervised consensus clustering analysis: distance = “ euclidean”, clusterAlg = “ km”, pItem = 0.8, pFeature = 1, seed = 2425, maxK = 10, reps = 100. Following consensus clustering analysis, further inspection of the consensus distribution (eCDFs) for each choice of K (i.e, the number of clusters) results in the identification of 3 stable clusters in the data (Supplementary Figure 1) labeled as: IE (Immune enriched), IE-ECM (Immune Enriched Dense Extracellular Matrix-like) and ID (Immune Dessert) subtypes. For more details on the process of unsupervised class discovery using consensus clustering analysis, please see the consensusClusterPlus R package documentation. To enable robust classification of individual samples into one of the three identified clusters in independent datasets, a naïve bayes classifier^34^ was trained on canine tumor data to predict its assigned cluster from its TME profile. We chose the naïve bayes classifier because it is relatively simple to train and interpret compared to other related machine learning approaches and easily scalable to larger datasets. For training, we only considered cell types whose relative abundances could be measured across all datasets analyzed in this study. For added robustness against outliers, we incorporated a standard data augmentation strategy of randomly injecting gaussian noise (mean: 0, standard deviation: 0.01) into the training data prior to feeding it as input to the naïve bayes classifier^65^.

### Cell type abundance estimation independently from single cell data

To estimate the proportion of each cell type in each tumor sample processed using single cell RNA Sequencing, we borrowed cell count data corresponding to each cell type from the supplementary material of the recently published single cell RNA Sequencing study^41^. The cell type proportions were then estimated by dividing the number of cells of a particular cell type from a sample with the total number of sequenced cells from that sample. Following cell type proportion calculation, all cell type proportions were standardized to z-scores for visualization in a heatmap (Supplementary Figure 7A,B)

### Differential expression and gene set enrichment analysis

Raw read count data and the inferred TME subtype of each sample from the discovery cohort was provided as input to edgeR^66^ (v3.40.2) using default parameter settings for subtype-specific differential expression analysis. For each TME subtype, all genes were ranked by their log fold change in expression estimated by edgeR. For a given subtype, genes ranked at the top of the list correspond to genes with high positive log fold change in expression associated with that subtype (i.e have relatively higher expression in that subtype vs other subtypes), whereas genes ranked at the bottom of the list correspond to genes with high negative log fold change in expression associated with that subtype (i.e have relatively lower expression in that subtype vs other subtypes). For each subtype, the ranked list of all genes was then provided as input to the standard Gene Set Enrichment Analysis (GSEA) pipeline implemented in clusterProfiler^67^ (v4.6.2) with the following parameters specified: (nPermSimple: 100000, minGSSize = 10, maxGSSize = 500) to estimate the relative enrichment of cancer hallmark pathways in each subtype. For each subtype, the top 5 enriched pathways ranked in decreasing order by the normalized enrichment score were selected for downstream biological interpretation.

### Survival analysis

To perform Kaplan-Meier and cox proportional hazards survival analysis of canine clinical trial and human clinical datasets, we utilized the survival (v3.5.5) and survminer (v0.4.9) packages implemented in R. In the case of canines, adjuvant sirolimus treatment did not affect any of the trial endpoints, inclusive of disease-free interval or overall survival. A post-hoc analysis of the pharmacokinetic parameters from dogs receiving standard of care + adjuvant sirolimus indicate a high degree of variability in exposures and a lower-than-expected trough blood level, below the predicted target of 10 uM^27^. Thus, these canine patients were considered together in the survival analyses presented herein (Figures 3 and 4). In the case of human Osteosarcomas, all 405 patients analyzed in this study received standard of care neoadjuvant chemotherapy, with majority of these patients (N=383) receiving a diagnostic biopsy of their primary tumor prior to neoadjuvant chemotherapy treatment. Hence these patients were considered together in downstream survival analysis. 185 out of these 383 patients had matched progression free survival outcome information measured, which was standardized to the time interval (in days) from the time of diagnosis to the time of relapse or metastasis. This information was utilized to perform Kaplan-Meier survival analysis and plot progression free survival trajectories of patients stratified by their TME subtype (Figure 5D). Of these 185 patients, 83 patients had information on percent tumor necrosis following histopathological evaluation after neoadjuvant chemotherapy treatment (>90%, <= 90%) along with age, gender and stage of disease at initial diagnosis (localized vs metastatic). This information was utilized to fit a cox proportional hazards regression model utilizing the following variables as features: age, gender, stage, percent necrosis and TME subtype (Figure 5E). For an independent cohort of patients (N=53 out of 84 patients from GSE33383^37^) a binary outcome indicating whether the patient progressed to metastatic disease within 5 years from initial diagnosis was measured. This information was utilized to estimate the proportion of patients developing metastasis within 5 years for each TME subtype (Figure 5F).

## Data availability

The canine clinical trial datasets generated during and/or analyzed during the current study are available from the corresponding author on reasonable request.

## Code availability

All the codes required to reproduce the results of this study are deposited at: https://github.com/spatkar94/doghuman_TME_deconvolution.git

## Acknowledgements

We would like to thank the CCR Sequencing Facility located at the Frederick National Laboratory for Cancer Research (FNLCR), NCI, NIH, Frederick, MD 21701, for initial data analysis and RNA Sequencing of canine tumor data.

