## Supplementary Figures for "Large Scale Comparative Deconvolution Analysis of the Canine and Human Osteosarcoma Tumor Microenvironment Uncovers Conserved Clinically Relevant Subtypes"

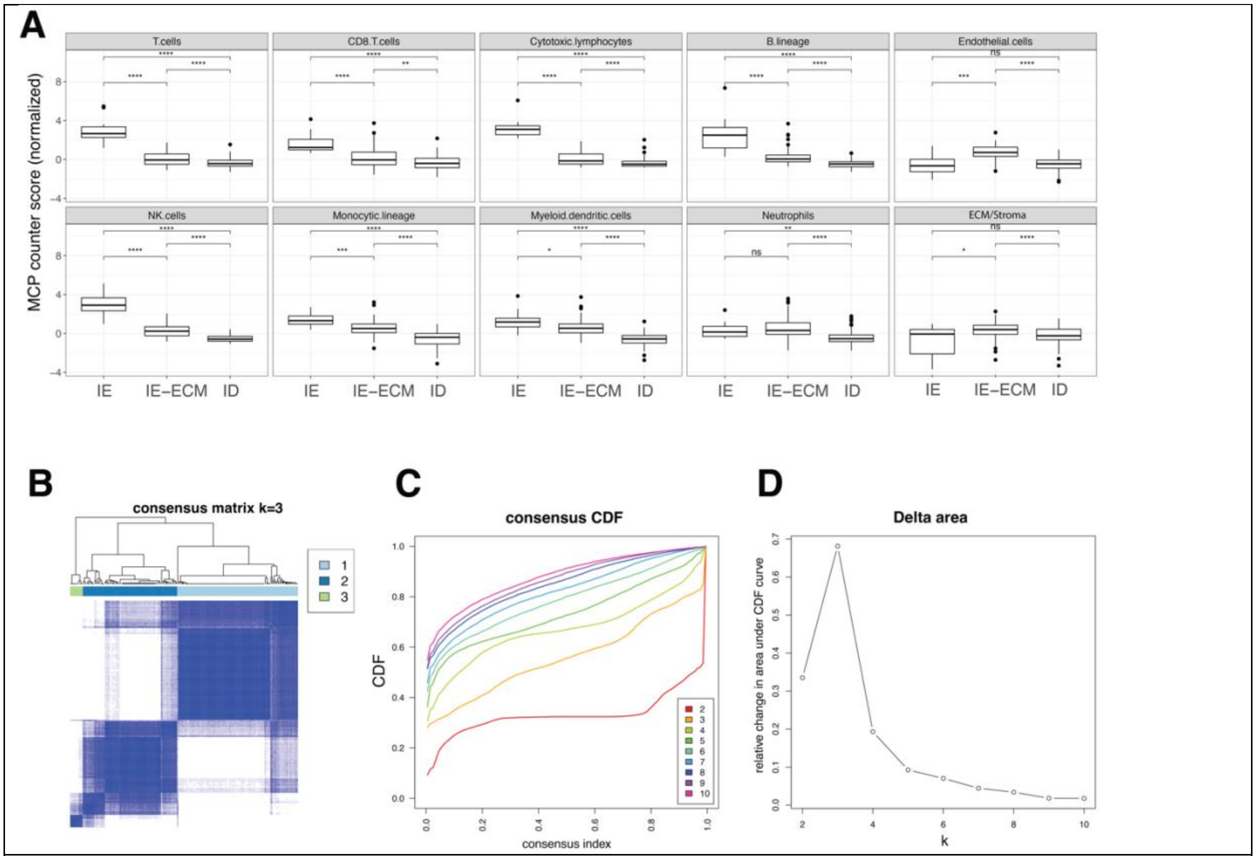


**Supplementary Figure 1 A:** Boxplots depicting the MCP counter-estimated relative cell type abundance of each cell type/cell lineage in the Discovery Cohort (N=198). Statistical significance between groups was determined by non-parametric Wilcoxon rank sum test (p-value < 0.05: *, p-value < 0.01: **, p-value < 0.001: ***, p-value < 0.0001: ****). **B:** Consensus matrix depicting the consensus distribution over all pairs of samples for N=3 clusters assigned by K-means clustering. **C:** Empirical CDF of consensus distribution. **D:** relative change in consensus as we increased number of clusters. At N=3 clusters we see the largest appreciable increase in consensus, thus suggesting the existence of 3 stable clusters in the data based on MCP counter-estimated cell type abundances

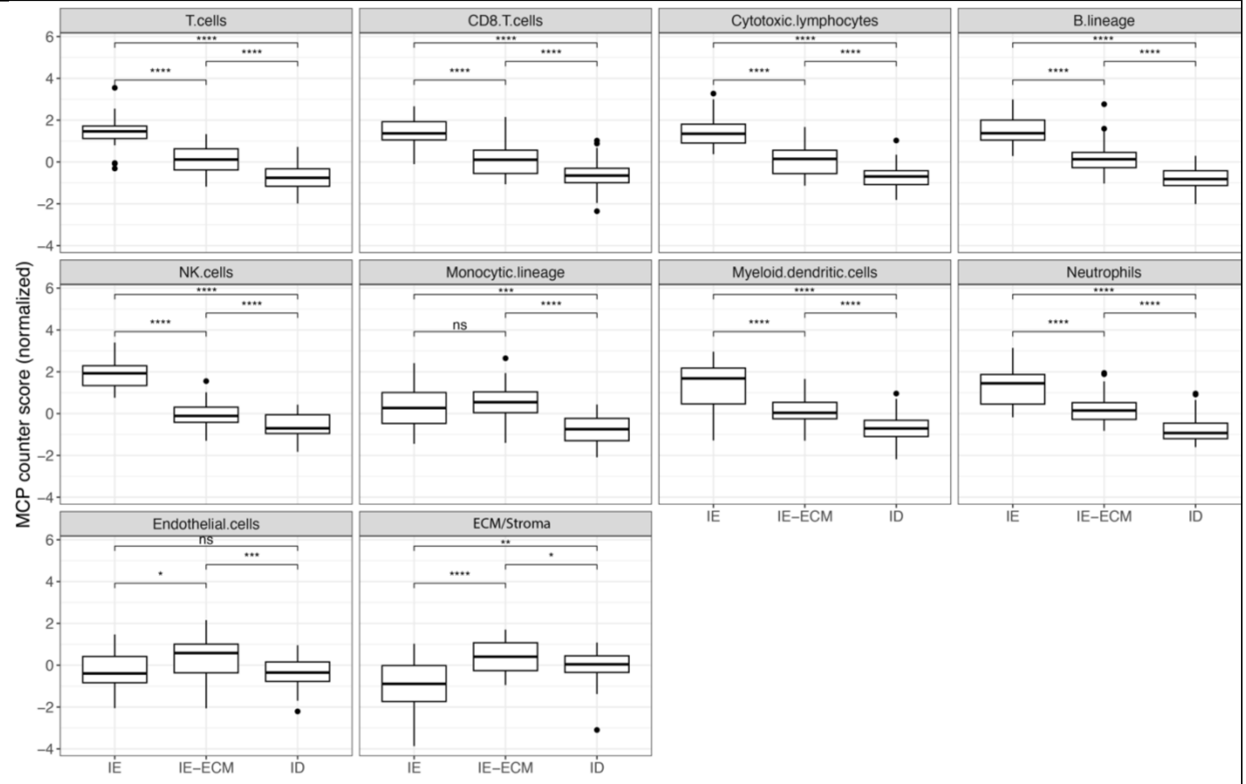


**Supplementary Figure 2.** Boxplots depicting the MCP counter-estimated relative cell type abundance of each cell type/cell lineage in the Validation Cohort (N=115 samples) and grouped by TME subtype. Statistical significance between groups was determined by non-parametric Wilcoxon rank sum test (p-value < 0.05: *, p-value < 0.01: **, p-value < 0.001: ***, p-value < 0.0001: ****).

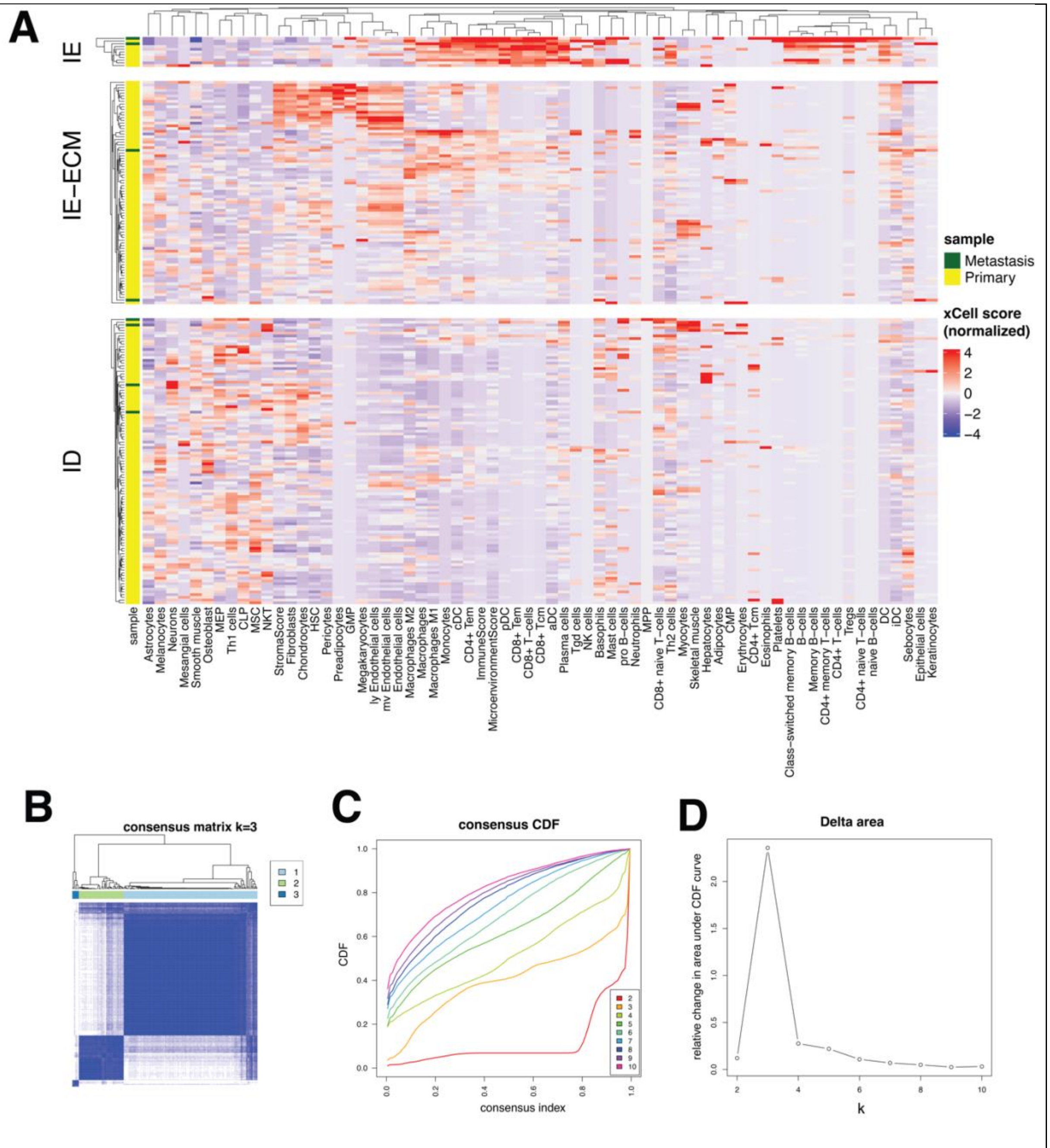


**Supplementary Figure 3 A:** Heatmap depicting the xCell-estimated relative cell type abundances in each sample for the discovery cohort (N=198 samples). **B:** Consensus matrix depicting the consensus distribution over all pairs of samples for N=3 clusters assigned by K-means clustering. **C:** Empirical CDF of consensus distribution. **D:** relative change in consensus as we increased number of clusters. At N=3 clusters we see the largest appreciable increase in consensus, thus suggesting the existence of 3 stable clusters in the data based on xCell-estimated cell type abundances.

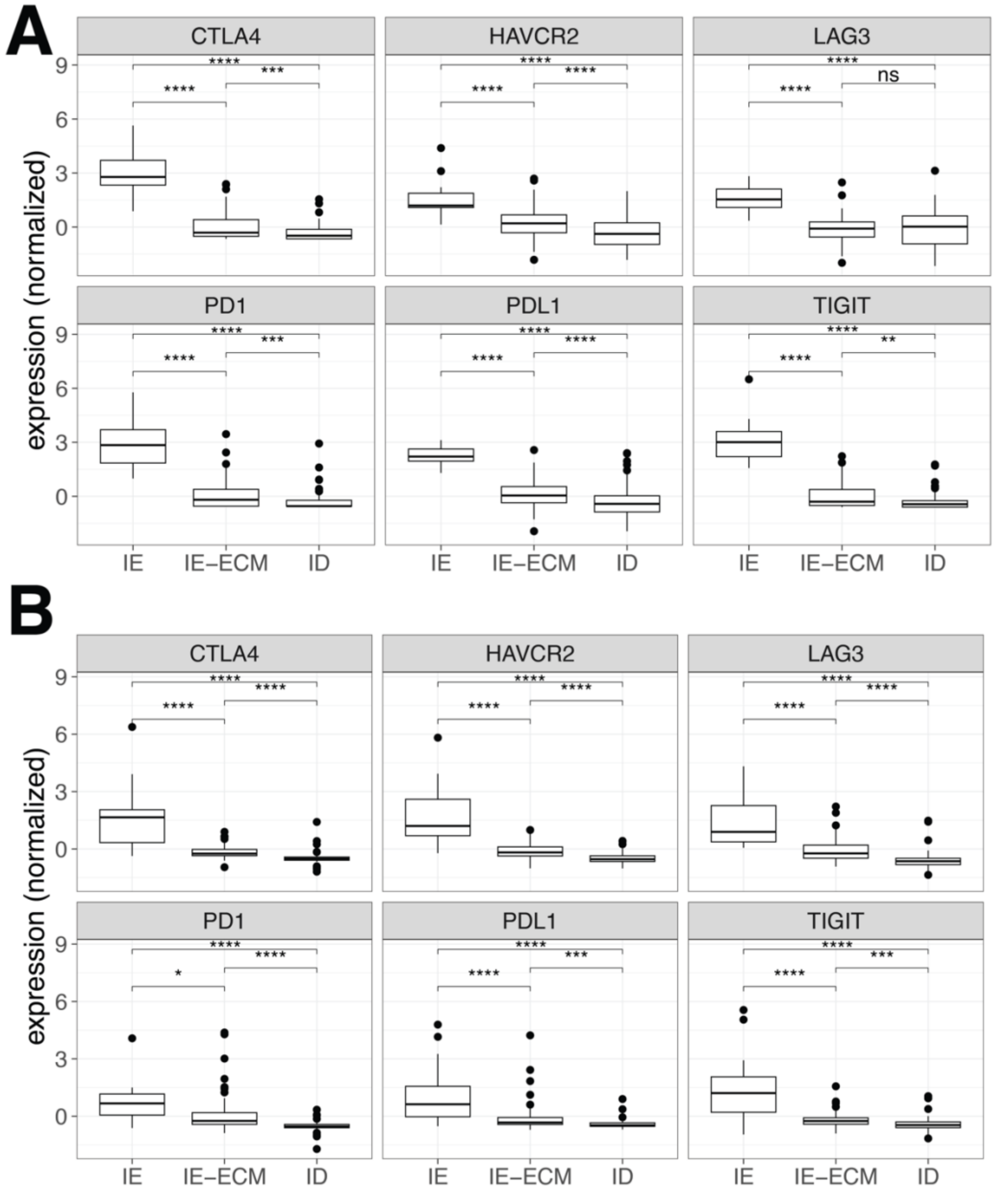


**Supplementary Figure 4 A-B:** Boxplots depicting the normalized expression of immune checkpoint genes in (A) the Discovery Cohort (N=198 samples) and (B) the validation cohort (N=115 samples) and grouped by TME subtype. Statistical significance between groups was determined by non-parametric comparison using Wilcoxon rank sum test (p-value < 0.05: *, p-value < 0.01: **, p-value < 0.001: ***, p-value < 0.0001: ****)

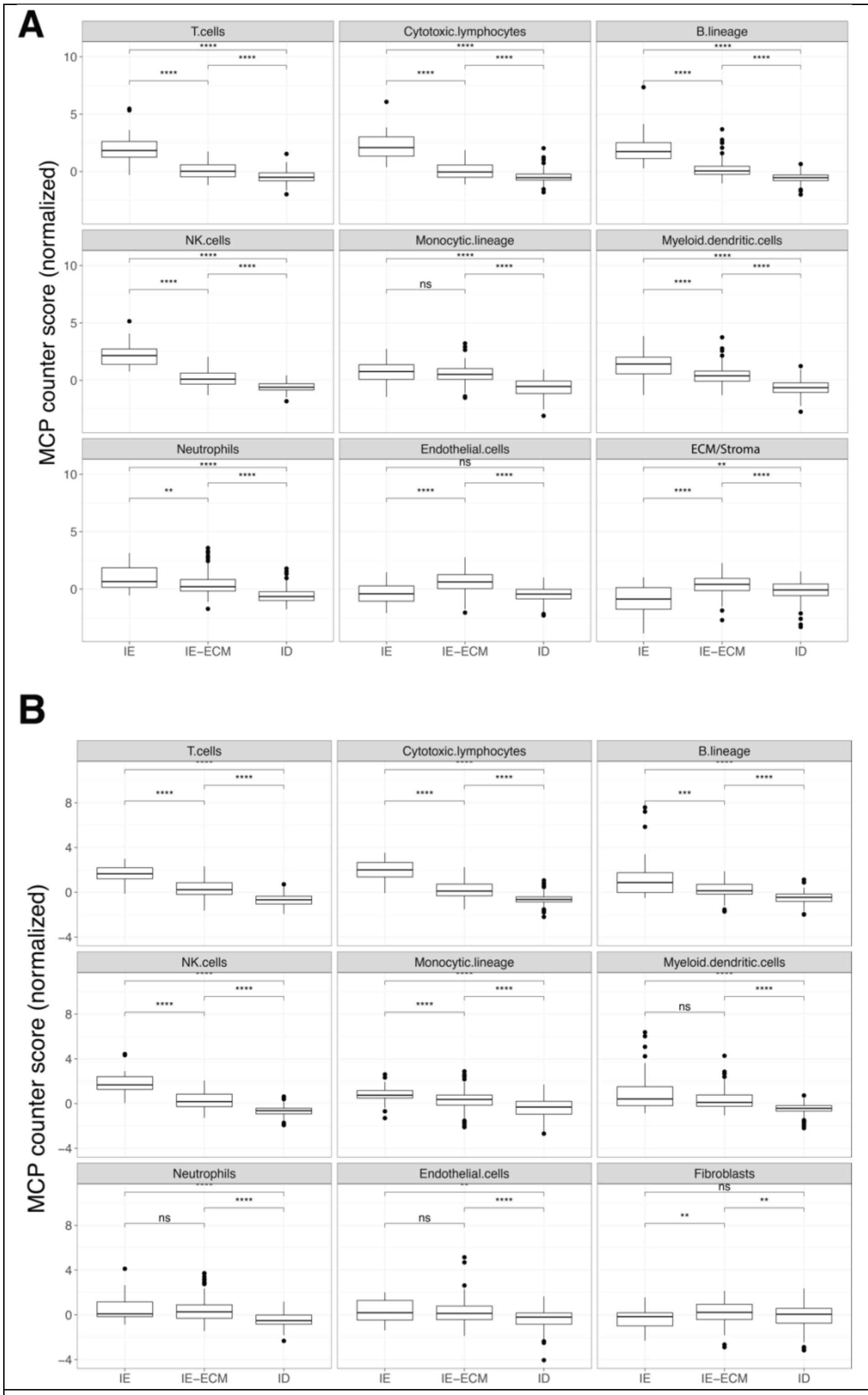


**Supplementary Figure 5. A-B:** Boxplots depicting the MCP counter-estimated relative cell type abundance of each cell type/cell lineage in (A) canine Osteosarcoma (Discovery + Validation cohort) and (B) human Osteosarcomas (N=405 samples) grouped by TME subtype. Statistical significance between groups was determined by non-parametric Wilcoxon rank sum test (p-value < 0.05: *, p-value < 0.01: **, p-value < 0.001: ***, p-value < 0.0001: ****)

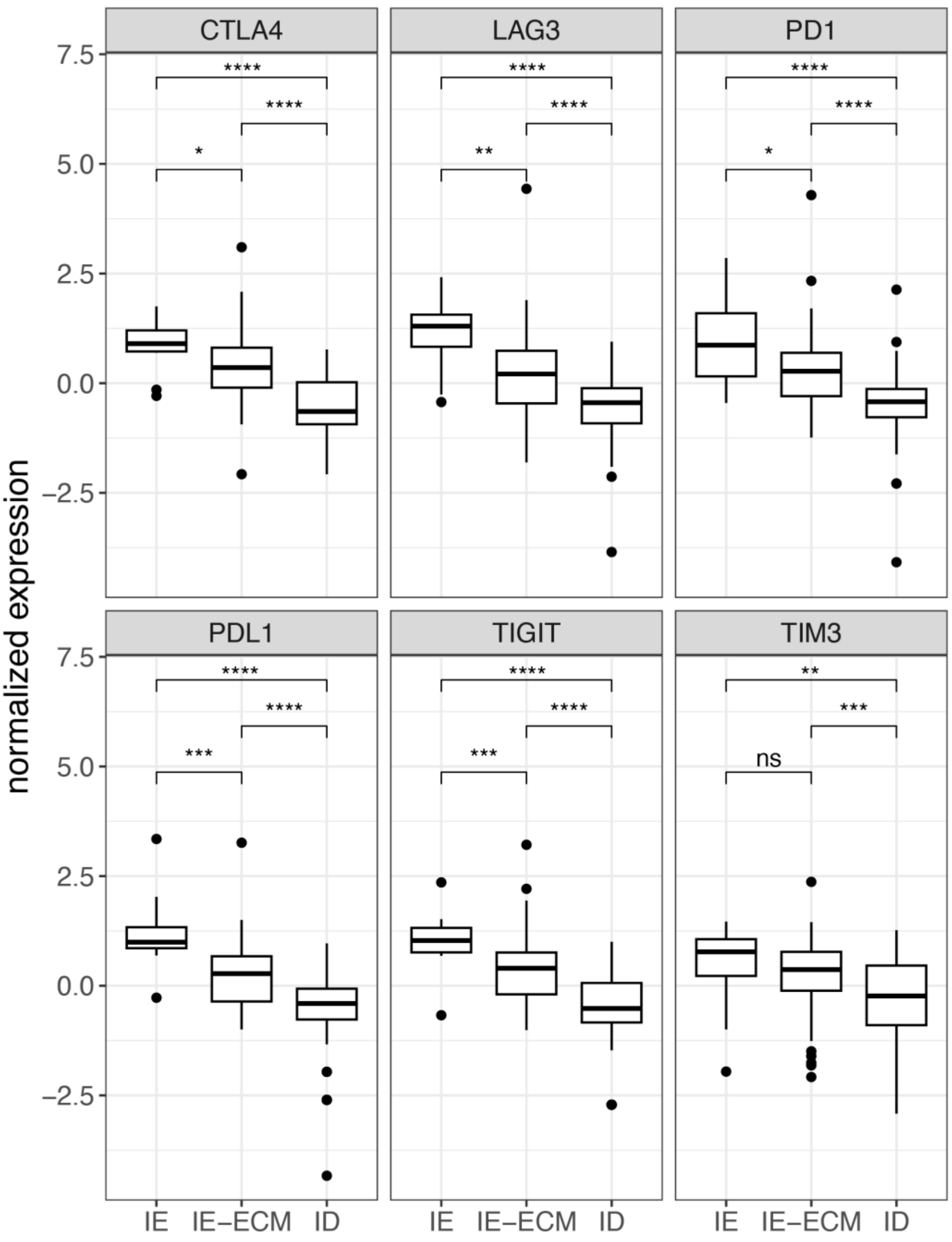


**Supplementary Figure 6.** Boxplots depicting the normalized expression of immune checkpoint genes in human Osteosarcomas (N=405 samples) and grouped by TME subtype. Statistical significance between groups was determined by non-parametric comparison using Wilcoxon rank sum test (p-value < 0.05: *, p-value < 0.01: **, p-value < 0.001: ***, p-value < 0.0001: ****)

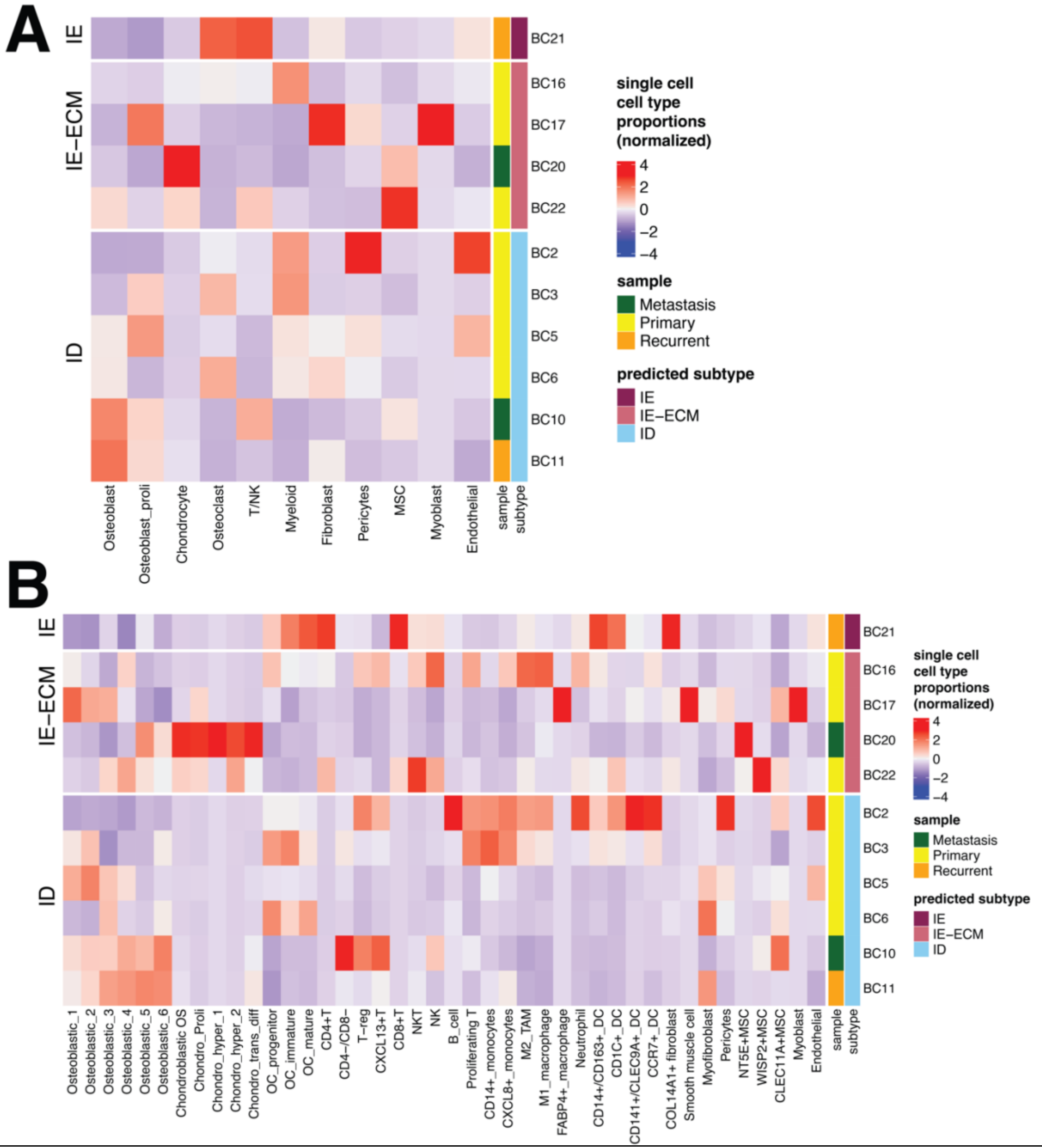


**Supplementary Figure 7 A-B:** Heatmap depicting the cell type proportions of distinct cell types (A) and their respective subtypes (B) in individual human Osteosarcoma tumor samples identified from Single Cell RNA Sequencing analysis (GSE152048).

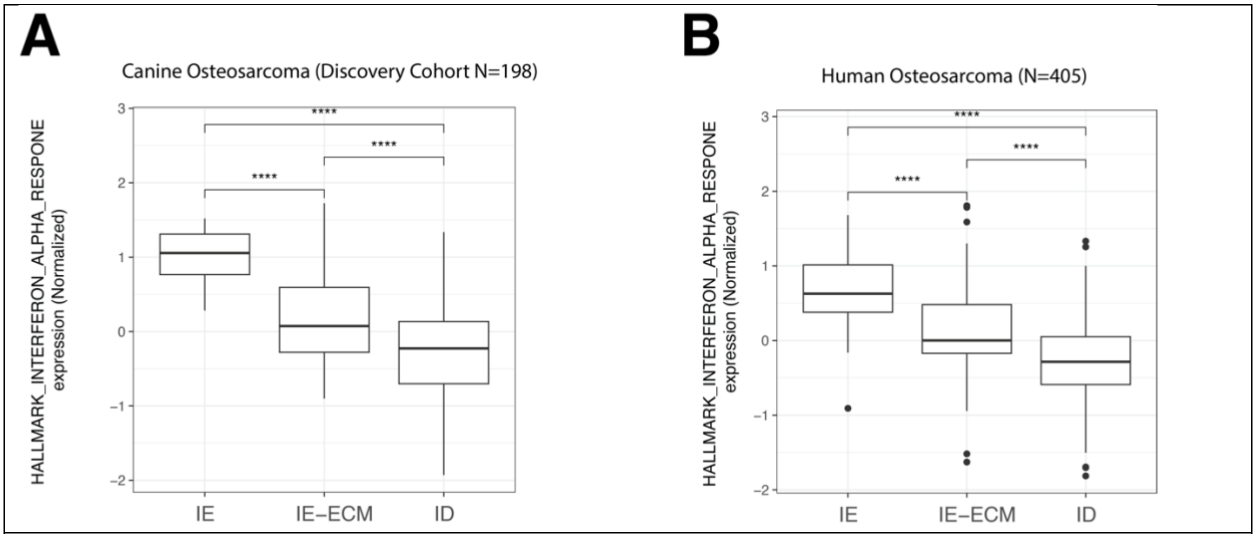


**Supplementary Figure 8 A-B:** Boxplots depicting the average normalized expression of genes part of Type 1 interferon response signaling cancer hallmark pathway in canine Osteosarcomas (A) and human Osteosarcomas (B)
